# SenSeqNet: A Deep Learning Framework for Cellular Senescence Detection from Protein Sequences

**DOI:** 10.1101/2024.10.28.620702

**Authors:** Hanli Jiang, Li Lin, Dongliang Deng, Jianyu Ren, Xin Yang, Siyi Liu, Lubin Liu

## Abstract

Cellular senescence, characterized by the irreversible cessation of division in normally proliferating cells due to various stressors, presents a significant challenge in the treatment of age-related diseases. Understanding and accurately detecting cellular senescence is crucial for identifying potential therapeutic targets. However, traditional wet lab assays for detecting cellular senescence are time-consuming and labor-intensive, limiting research and drug development efficiency. There is an urgent need for computational tools allowing swift and accurate detection of cellular senescence from protein sequences. We propose SenSeqNet, a novel deep learning framework for detecting cellular senescence directly from protein sequences. The framework begins with feature extraction using the Evolutionarily Scaled Model (ESM-2), a state-of-the-art protein language model that captures evolutionary information and complex sequence patterns. The extracted embeddings are then passed through a hybrid architecture consisting of long short-term memory (LSTM) networks and convolutional neural networks (CNNs) to further refine and learn from the embedded information. SenSeqNet achieved a final accuracy of 83.55% on independent testing, surpassing various machine learning and deep learning architectures. This performance underscoring the robustness and effectiveness of SenSeqNet for detecting cellular senescence from protein sequences. These results provide a solid foundation for future research on aging and age-related therapeutics.

## 1. Introduction

Aging is a complex biological process marked by the gradual accumulation of cellular and molecular damage over time, leading to diminished physiological function and the onset of age-related diseases^1^. The incidence rates of diabetes, heart disease, neurological disorders, and various cancers increase with age^2^. Therefore, the early prediction and detection of cellular senescence play a vital role in preventing the development of numerous diseases^3^. Carlos López-Otín et al. have identified loss of proteostasis as a key hallmark of aging^4^. Recent advances in high-throughput sequencing and proteomics have generated an unprecedented volume of biological data, presenting new opportunities to explore aging at the molecular level. Protein sequences, in particular, are of significant interest because they play a direct role in nearly all cellular processes. Analyzing these sequences to identify signs of aging could reveal key biomarkers and therapeutic targets, paving the way for novel interventions in age-related conditions.

Traditional methods for studying protein sequences have often relied on manual analysis and rule-based approaches, which are limited in their capacity to uncover the complex patterns associated with aging^5^. However, the emergence of machine learning (ML) and deep learning (DL) has revolutionized the analysis of biological data, enabling the automatic extraction of complex patterns and insights that were previously unattainable with traditional methods^6,7^. In particular, deep learning models, with their ability to automatically learn hierarchical representations from raw data, have demonstrated remarkable success in various biomedical applications, including protein structure prediction, drug discovery, and disease diagnosis^8-10^.

Despite significant advancements in deep learning, the application of these techniques to detect cell aging using protein sequences remains relatively scarce. Recently, the integration of multiple deep learning models to leverage their respective advantages has demonstrated significant success^11^. To further explore the application of integrated deep learning models, we developed SenSeqNet (Fig. 1) , a novel deep learning approach that begins with ESM2 for feature extraction and then synergistically combines the strengths of long short-term memory (LSTM) networks and convolutional neural networks (CNNs)^12^. LSTM networks are adept at modeling sequential data like protein sequences, capturing long-term dependencies that are critical for understanding the temporal aspects of aging^13^. Meanwhile, CNNs excel at extracting spatial features^14^, making them ideal for identifying complex patterns within the outputs generated by LSTM networks.

**Fig. 1.**
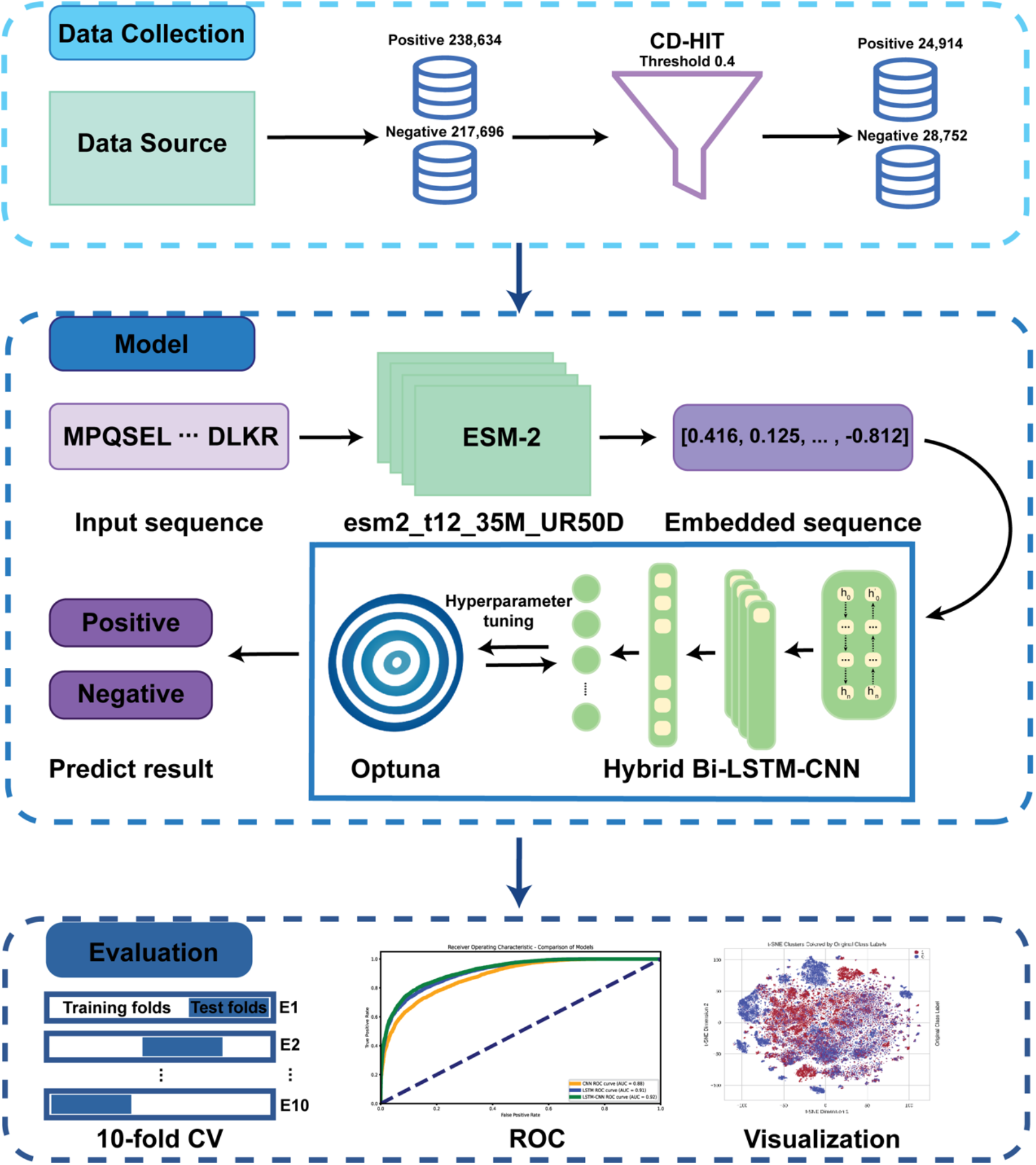
The overall framework.

The next crucial step in sequence-based protein prediction tasks is choosing the optimal model architecture, with the representation of sequence data being a key factor in determining overall model performance^15^. We utilized the Evolutionarily Scaled Model (ESM)^16^, a state-of-the-art transformer-based model renowned for its ability to capture deep evolutionary relationships in protein sequences. Specifically, we evaluated several variants of the ESM-2 model to generate embeddings for our task. To gain insights into the high-dimensional data structure, we conducted visualization analysis using the t-distributed Stochastic Neighbor Embedding (t-SNE) algorithm^17^. This allowed us to observe distinct clustering patterns within the protein sequence embeddings, highlighting the effectiveness of our embedding approach. While other PLMs such as ProtTrans ^18^, ESM-1b^19^, ESM-1v^20^ and ProteinBERT^21^ were considered, their sequence length limitations (e.g., 1024 amino acids) made them less suitable for our dataset, which contains nearly 4000 protein sequences exceeding this limit. ESM-2, with its support for sequences exceeding 1024 amino acids, was chosen for its ability to handle longer sequences while preserving global context, resulting in consistent and robust performance in preliminary experiments. We also compared the t-SNE visualizations between ESM-2 and other traditional feature extraction methods. ESM-2 outperforms other approaches in producing more clearly defined and separable clusters. Through extensive experimentation, we evaluated multiple high-performance machine learning and deep learning models to identify the optimal architecture for classifying cell aging. We compared these models based on their ability to capture unique patterns in protein sequences related to cellular senescence. Ultimately, SenSeqNet demonstrated superior performance through independent testing, leveraging its ability to effectively process both sequential and spatial features. In addition to 10-fold cross-validation, external validation further confirmed that SenSeqNet is a highly robust tool for this task. This study advances the application of deep learning in aging research, providing a practical solution for large-scale screening of senescence-associated proteins and establishing a foundation for identifying potential biomarkers and supporting the development of targeted therapies to promote healthy aging.

## 2. Materials and methods

### 2.1. Benchmark datasets

For the development of an effective prediction model, the dataset is generally composed of comparable positive and negative samples. In this study, cell senescence genes were downloaded from the CellAge: The Database of Cell Senescence Genes, where abnormal expression of these genes is associated with the induction of senescence. The database can be accessed at https://genomics.senescence.info/cells/index.html. Protein sequences related to cell senescence were downloaded from the Universal Protein Knowledgebase (UniProtKB)^22^ and used as positive samples. In addition, protein sequences from housekeeping genes were selected as negative samples, resulting in a dataset comprising 238,634 positive and 217,696 negative protein sequences. Housekeeping genes are essential for basic cell maintenance, maintaining constant expression levels across all cells and conditions, and play a crucial role in calibration for various biotechnological applications and genomic studies^23^. To reduce redundancy and ensure high-quality data, we employed CD-HIT^24^ with a threshold of 0.4, ultimately leaving 24,914 positive and 28,752 negative sequences. The dataset will be divided into a training set (80%) and a validation set (20%) to facilitate model development and performance evaluation. The code and benchmark dataset are publicly available at https://github.com/HanliJiang13/SenSeqNet, ensuring transparency and reproducibility of the study.

### 2.2. SenSeqNet model structure

As illustrated in Figure 2, SenSeqNet consists with two main components: ESM-2 and the hybrid LSTM-CNN architecture. Beginning with the first component, effective representation of sequence data is an essential step in building a robust prediction model, especially when dealing with protein sequences^25^. Thus, we employed the ESM-2^16^, known for its exceptional ability to capture deep evolutionary relationships in protein sequences (Fig. 2A). ESM-2 is trained on approximately 65 million unique protein sequences using a masked language modeling objective, which involves predicting the identity of randomly selected amino acids within a sequence based on their surrounding context^14^. This training enables ESM-2 to learn the dependencies between amino acids, allowing it to effectively capture both local and global contextual information.

**Fig. 2.**
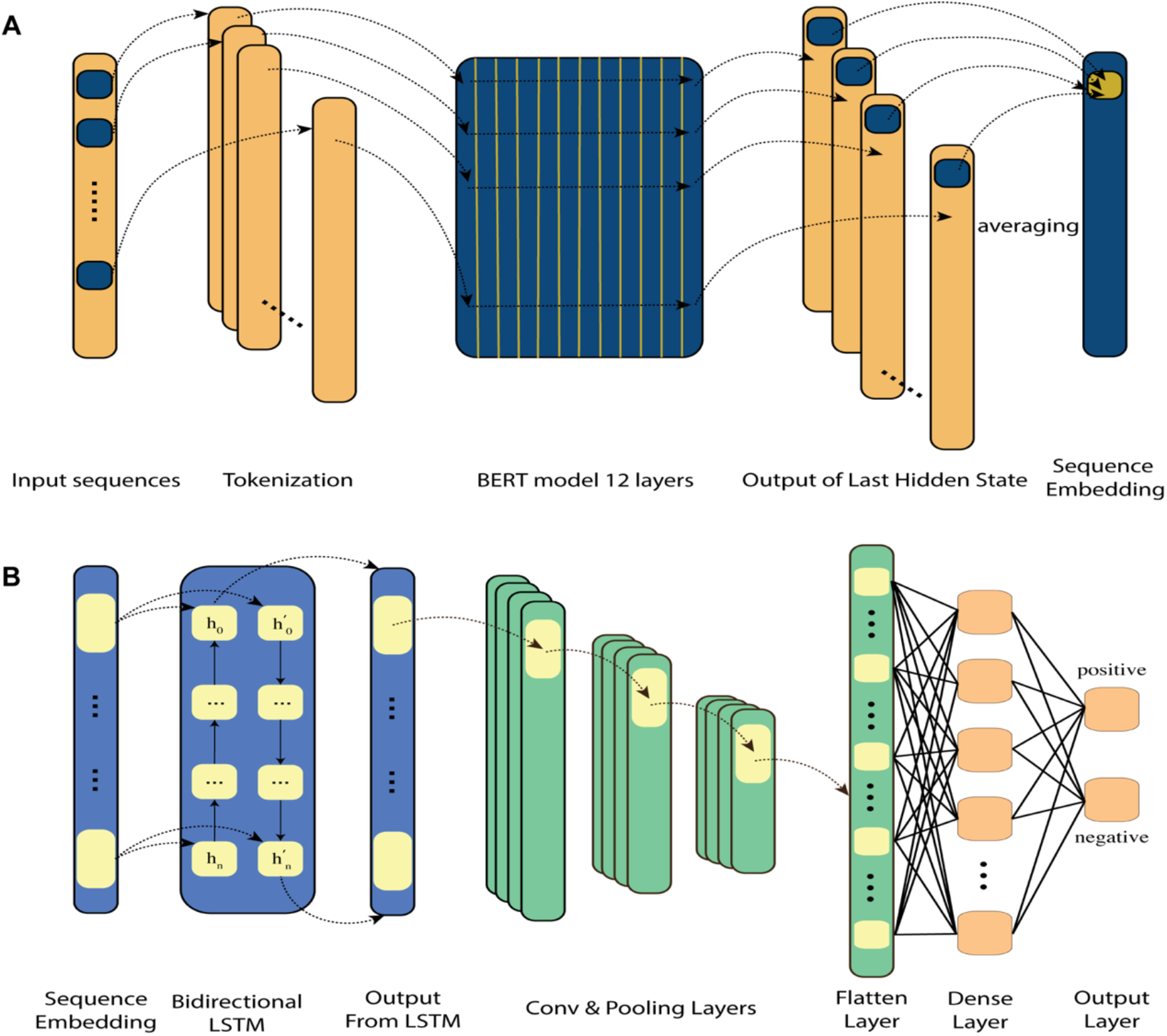
Sample Architecture of the SenSeqNet: A. Evolutionarily Scaled Model (ESM-2) architecture for feature extraction. B. Hybrid Long Short-Term Memory-Convolutional Neural Network (LSTM-CNN) model.

The unsupervised nature of the ESM-2 model’s training allows it to internalize sequence patterns across diverse proteins, effectively capturing structural information linked to these patterns^26^. ESM-2 is highly effective at processing and embedding from large-scale protein datasets, making it particularly suitable for sequence representation tasks. The model’s performance across various bioinformatics applications has been well-documented^27^, demonstrating its ability to discern subtle patterns within sequences, which are important for accurate prediction tasks. There are multiple ESM-2 models, each differing in architectural configuration, where an increase in the number of transformer layers corresponds to a proportional increase in the number of model parameters. To identify the most effective variant for our dataset, we evaluated each variant’s output and utilized the results to train two distinct models: a CNN to assess the spatial features and an LSTM to analyze the sequential dependencies within the results. By balancing the accuracy, computational efficiency, and scalability of these models, we determined the most effective ESM-2 variant for our dataset. The selected model variant was then used for embedding generation throughout our study. The detailed results of this evaluation will be discussed in the Results section.

Followed by ESM-2, the embedded sequences are input into the second main component, a hybrid LSTM-CNN architecture (Fig. 2B). In this setup, a Bidirectional LSTM network is first employed to process the sequence data, leveraging its capacity to model temporal dependencies in both directions, thus capturing comprehensive long-term relationships between amino acids in the protein sequences. These sequential features are then passed to the CNN, which focuses on identifying spatial patterns within the LSTM-extracted features. This combined approach enables the model to capture both temporal and spatial characteristics. This two-part architecture, combining the evolutionary context captured by ESM-2 with the temporal and spatial analysis capabilities of the LSTM-CNN, enables SenSeqNet to effectively identify crucial features in protein sequences. This comprehensive approach significantly enhances the model’s ability to distinguish aging cells from non-aging cells, as demonstrated in the results.

### 2.3. Comparative models for benchmarking SenSeqNet

In this study, we explored several machine learning and deep learning architectures to develop a reliable model for detecting cell aging using protein sequences^25^. Given the lack of prior research in this specific domain of cell aging, our approach involved comprehensive experimentation with a range of models commonly employed in protein sequence prediction to understand their capabilities and limitations. The architectures evaluated include CNNs^28^, Recurrent Neural Networks (RNNs)^29^, LSTMs^13^, and bidirectional LSTMs. Recognizing the strengths of both CNNs in extracting spatial features and bidirectional LSTMs in capturing sequential dependencies^30^, we also experimented with hybrid models that combined these two architectures. The combined CNN and bidirectional LSTM model was designed to leverage the advantages of both, aiming to improve the accuracy and robustness of cell aging detection from protein sequences. Additionally, we constructed several machine learning models, including Bagging^31^, Support Vector Machines (SVM)^32^, Random Forest (RF)^33^, and XGBoost^34^, to provide a comparative understanding of their performance against deep learning approaches. Below, we present a comprehensive description of each deep learning model. Hyperparameters such as learning rate, dropout rate, batch size, and the number of layers were optimized for each model using Optuna^35^, a framework for hyperparameter optimization.

#### 2.3.1. Convolutional Neural Network

This study introduces a custom CNN Classifier for sequence prediction, employing a multi-layer convolutional neural network architecture. The model consists of three convolutional layers, each with a varying number of filters (79, 111, and 180 filters, respectively) to extract spatial features from the input sequences. Max-pooling layers follow the first two convolutional layers to reduce the spatial dimensions of the feature maps, and batch normalization is applied after each convolutional layer to stabilize training. The flattened feature maps are then passed into fully connected layers consisting of 512 and 256 neurons to learn higher-level feature representations. Dropout is applied between the fully connected layers to prevent overfitting, and the final output layer produces multi-class predictions using a log softmax activation function. This model efficiently captures both local and global feature representations through the combination of convolutional and fully connected layers.

#### 2.3.2. Recurrent Neural Networks

This study introduces a custom RNN Classifier for sequence prediction, utilizing a multi-layer RNN architecture. The model incorporates an RNN with multiple layers to capture temporal dependencies in the input sequences. The hidden layers of the RNN consist of a specified number of units, optimized for extracting sequential features. A batch normalization layer is applied to the output of the last time step to stabilize training and improve convergence. The model’s sequential output is further passed through fully connected layers consisting of 512 and 256 neurons, respectively, to capture higher-level feature representations. Dropout is applied between the fully connected layers to reduce overfitting. This model efficiently captures both temporal patterns and higher-order representations through its recurrent and fully connected layers.

#### 2.3.3. Long Short-Term Memory Network

The LSTM Classifier we used, incorporates LSTM layers that are designed to capture long-term dependencies in the input sequences by considering both past and future information at each time step. Specifically, the model first processes input sequences through a stacked LSTM layer containing 427 LSTM units, allowing it to retain relevant information across time steps. The LSTM is set with multiple layers (num_layers = 5) to improve its capacity to capture complex temporal relationships. The output from the final LSTM layer, representing the last time step, is passed through a dropout layer to mitigate overfitting. This is followed by a fully connected layer, which transforms the LSTM output into multi-class predictions. The model effectively leverages the sequential nature of LSTMs.

#### 2.3.4. Bidirectional LSTM

The Bi-LSTM Classifier extends the capabilities of traditional LSTMs by incorporating bidirectional LSTM layers. These layers are uniquely designed to process information both forwards and backwards along the input sequence, enabling the model to capture context from both the past and future at each time step. This dual-direction processing allows for a more comprehensive understanding of temporal relationships within the data. The Bi-LSTM architecture features a bidirectional LSTM layer with 250 hidden units distributed across 4 layers, ensuring that each time step is enriched with full sequence context. Such an arrangement not only enhances the model’s ability to discern patterns that a single-direction LSTM might miss but also improves predictive accuracy by integrating insights from the entire sequence breadth.

#### 2.3.5. Combined architectures

CNNs are particularly adept at capturing spatial information, making them highly effective for tasks that involve the analysis of grid-like data structures, such as images or sequences with local spatial dependencies. Additionally, LSTMs excel in handling sequential data, allowing for the modeling of long-term dependencies and temporal patterns, which are crucial in many predictive tasks. The integration of these two models aims to create a model that synergizes the spatial feature extraction capabilities of CNNs with the sequence modeling strengths of LSTMs. To explore the potential of this combination, we investigated several architectural strategies:

a) **CNN followed by LSTM (CNN-LSTM):** This architecture begins with the CNN, which extracts spatial features from the raw protein sequences. The CNN is adept at identifying local patterns or motifs within the sequence data that may be associated with cell aging. These spatial features are then fed into the Bidirectional LSTM, which captures the sequential dependencies among the CNN-extracted features. The Bidirectional LSTM’s ability to analyze the sequence data from both directions ensures that critical temporal information is not overlooked.
b) **Parallel LSTM and CNN (pLSTM-CNN)**: In this approach, the Bidirectional LSTM and CNN operate simultaneously on the input data. The Bidirectional LSTM processes the sequence data to capture temporal dependencies from both past and future directions, while the CNN extracts spatial features independently. The outputs of both models are then combined to form a comprehensive representation of the protein sequences, which is used for the final classification. This parallel architecture is designed to maximize the strengths of both models, capturing a wide range of information from the data by considering both spatial and sequential aspects simultaneously.

### 2.4. Evaluation Under Imbalanced Dataset Conditions

While our primary dataset was composed of an equal number of senescence-associated (positive) and non-senescent (negative) proteins (1:1 ratio), real-world distributions are often skewed^36^. To investigate how SenSeqNet performs under more realistic class imbalances, we constructed additional datasets with 1:2, 1:3, 1:5, and 1:10 ratios of positive to negative samples. Specifically, we randomly sampled a subset of positive proteins to achieve the desired ratios, while retaining all negative sequences. The resulting datasets were shuffled and split into training and testing subsets using the same procedures as in the balanced case.

Importantly, no further fine-tuning or hyperparameter adjustments were applied to our model. We used the same network architecture, learning rate, and training epochs across all imbalanced datasets. This decision reflects our goal of applying SenSeqNet as a broad screening tool, wherein minimal re-training efforts are preferred for rapid deployment on large-scale proteomic datasets. By assessing these imbalanced scenarios under the same training pipeline, we aimed to characterize how performance naturally shifts without additional model-specific optimization.

### 2.5. Evaluation metrics

The following commonly used evaluation metrics were employed to assess the effectiveness of the models^36^: sensitivity (Sn), specificity (Sp), Matthews correlation coefficient (MCC)^37^, accuracy (Acc), F1 score, and AUC. The formulas for these metrics are provided below:

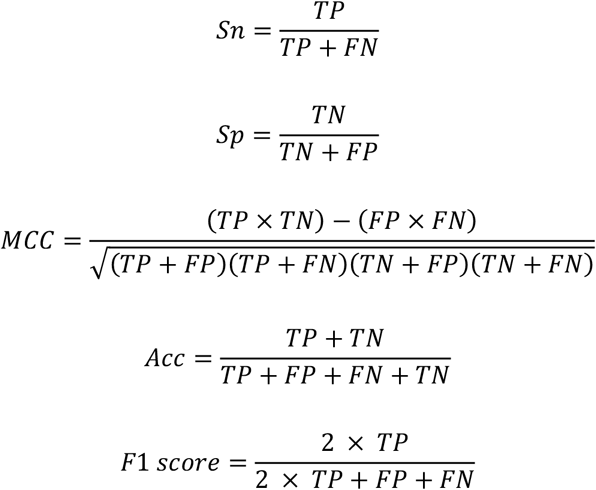

Where:

1) True Positive (TP): The number of correctly identified cases of cell aging.
2) True Negative (TN): The number of correctly identified non-aging cases.
3) False Positive (FP): The number of non-aging cases incorrectly identified as aging.
4) False Negative (FN): The number of aging cases incorrectly identified as non-aging.

AUC represents the area under the receiver operating characteristic (ROC) curve and is used to evaluate the overall performance of the model across different thresholds.

## 3. Results and discussion

### 3.1. ESM model selection

As detailed in the Methods section, we evaluated several variants of the Evolutionarily Scaled Model (ESM) for feature extraction from our protein sequence dataset. The goal was to identify the model that would provide the most accurate features while maintaining computational efficiency. Here, we present the results of this evaluation and the rationale for selecting the optimal ESM variant. The ESM variants evaluated included ESM-2_t6, ESM-2_t12, ESM-2_t30, and ESM-2_t33, each varying in complexity and embedding dimensions. The performance of these models was assessed based on their average accuracy across both the CNN and LSTM models, as well as the time required for feature extraction(Table 1).

**Table 1:** Comparison of ESM-2 Model Variants for Feature Extraction.

| Model | #Layers | #Params(M) | #Emb Dim | Acc(%) | Time |
| --- | --- | --- | --- | --- | --- |
| ESM-2_t6 | 6 | 8 | 320 | 79.50 | 1 |
| ESM-2_t12 | 12 | 35 | 480 | **81.37** | 2 |
| ESM-2_t30 | 30 | 150 | 640 | 78.71 | 4 |
| ESM-2_t33 | 33 | 650 | 1280 | 79.93 | 7 |

*Note: 1. Time: The "Time" column represents the relative feature extraction time for each ESM-2 model variant. The extraction time for ESM-2\_t6 is set as the baseline (1), and the times for other models are scaled relative to it. For example, ESM-2\_t12 takes twice as long as ESM-2\_t6, ESM-2\_t30 takes four times as long, and ESM-2\_t33 takes seven times as long.*

The ESM-2_t12 model achieved the highest accuracy at 81.37%, outperforming the other variants. This accuracy advantage, coupled with its moderate computational demands, makes ESM-2_t12 the most effective choice for feature extraction. Therefore, we have selected ESM-2_t12 for use throughout our experiments. To provide a more intuitive illustration of how ESM-2_t12 embeddings differentiate the classes, we performed t-SNE on the extracted features (Fig. 3A). In contrast to the manual feature extraction methods—ACC (Fig. 3B), CKSAAP (Fig. 3C), and DDE (Fig. 3D)— which exhibit heavily overlapping clusters, the ESM-2_t12 embeddings form distinct groupings that closely align with the original class labels. This clear separation indicates that ESM-2_t12 captures critical sequence-level signals pertinent to senescence, rather than arbitrary noise. Moreover, the t-SNE plots offer a complementary perspective to numerical metrics such as accuracy, specificity, and MCC, thereby reinforcing our confidence that ESM-2_t12 provides a robust foundation for downstream classification tasks.

**Fig. 3.**
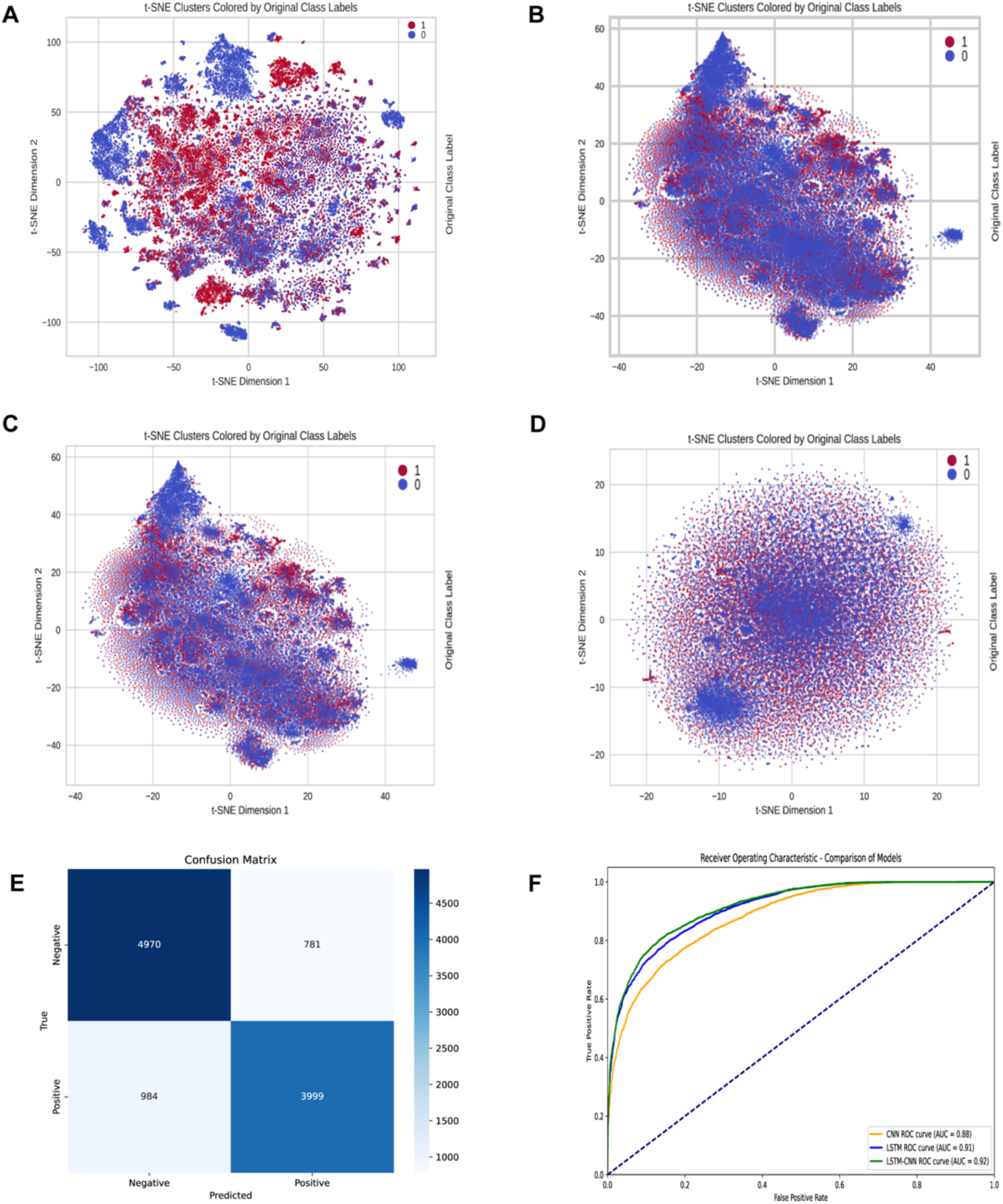
t-SNE visualization for ESM-2 and traditional methods A-D: A. t-SNE visualization for the ESM-2 model. B. t-SNE visualizetion for the ACC. C. t-SNE visualizetion for the CKSAAP. D. t-SNE visualizetion for the DDE. E. Confusion matrix for the SenSeqNet, illustrating a balanced classification outcome. F. Receiver Operating Characteristic (ROC) curves for the CNN, LSTM, and SenSeqNet models, comparing true positive rates across various thresholds. The curves highlight the superior sensitivity and specificity of SenSeqNet, which effectively combines the strengths of both CNN and LSTM architectures.

### 3.2. Comparative Performance of Multiple Protein Language Models

To assess the generalizability of SenSeqNet, we extracted features from multiple protein language models (PLMs), including ProtTrans (T5), ProteinBERT, ESM-1b, ESM-1v, and ESM-2, and subsequently fed these embeddings into our LSTM-CNN architecture. Table 2 presents the performance metrics across all tested configurations.

**Table 2:** Comparative performance of using embeddings from ProtTrans, ProteinBERT, ESM-1b, and ESM-2.

| PLM | Acc% | Sn% | Sp% | F1-Score% | MCC |
| --- | --- | --- | --- | --- | --- |
| ProtTrans (T5) | 51.54 | 31.70 | 68.71 | 37.77 | 0.0044 |
| ProteinBERT | 60.94 | 41.83 | 77.48 | 49.84 | 0.2073 |
| ESM-1b | **84.57** | 83.86 | 85.47 | **85.92** | **0.6898** |
| ESM-1v | 83.48 | **84.53** | 82.27 | 84.52 | 0.6680 |
| ESM-2 | 83.55 | 79.53 | **87.03** | 81.92 | 0.6689 |

Our results reveal a stark contrast between the ESM series and the other PLMs evaluated. ESM-1b achieved the highest accuracy (84.57%), with strong sensitivity (83.86%), specificity (85.47%), and an F1-score of 85.92%. ESM-1v and ESM-2 also exhibited robust performance, attaining accuracies of 83.48% and 83.55%, respectively. These findings suggest that ESM-based embeddings capture key senescence-related patterns in protein sequences, even when considering longer sequences exceeding 1,024 residues.

Although ESM-1b achieved slightly higher accuracy than ESM-2 for this specific task, ESM-2 was selected for several key reasons. First, scalability and efficiency: ESM-2 processes longer sequences (over 1024 residues) without requiring extensive chunking and merging, and feature extraction with ESM-1b and ESM-1v required approximately five times more computational time compared to ESM-2. Second, active development: As the most recent iteration of the ESM family, ESM-2 benefits from ongoing updates, improving scalability and offering potential for further accuracy gains. Third, comparable high accuracy: While ESM-1b achieved slightly higher accuracy (84.57% vs. 83.55%), ESM-2 demonstrated comparable performance, with superior specificity (87.03%) and a robust Matthews correlation coefficient (MCC) of 0.6689, ensuring reliable predictions.

In contrast, ProtTrans (T5) and ProteinBERT, even when applying standard chunking and averaging strategies to handle sequences exceeding 1024 residues, performed significantly worse than ESM-based models. ProtTrans (T5) achieved only 51.54% accuracy, 31.70% sensitivity, 68.71% specificity, an F1-score of 37.77%, and an MCC of 0.0044, reflecting near-random classification. ProteinBERT performed moderately better, with 60.94% accuracy, but still fell far short of the performance achieved by ESM-based embeddings.

These results underscore the challenges faced by more generalized protein language models in distinguishing senescence-associated proteins from housekeeping proteins. In contrast, ESM-2 provided robust performance, computational efficiency, and scalability, making it the optimal choice for SenSeqNet and well-suited for large-scale analyses of aging-related proteins.

### 3.3. Comparative performance evaluation of SenSeqNet with other deep learning models

We compared the performance of SenSeqNet with several deep learning models, including CNN, RNN, LSTM, Bidirectional LSTMs and various hybrid models. Each model was tested to determine its effectiveness in detecting cell aging from protein sequences, focusing on key performance metrics such as accuracy, AUC, Sn, Sp, and MCC using an independent test est. The comparative results of these models are summarized in Table 3.

**Table 3:**
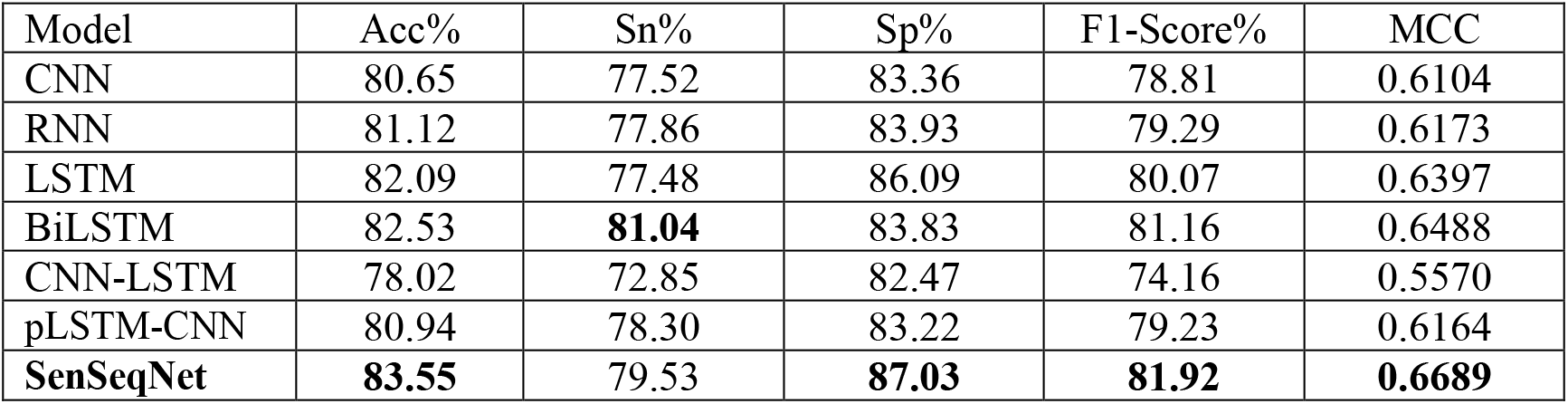
The performance comparison of SenSeqNet with other deep learning models on the independent test set.

| Model | Acc% | Sn% | Sp% | F1-Score% | MCC |
| --- | --- | --- | --- | --- | --- |
| CNN | 80.65 | 77.52 | 83.36 | 78.81 | 0.6104 |
| RNN | 81.12 | 77.86 | 83.93 | 79.29 | 0.6173 |
| LSTM | 82.09 | 77.48 | 86.09 | 80.07 | 0.6397 |
| BiLSTM | 82.53 | **81.04** | 83.83 | 81.16 | 0.6488 |
| CNN-LSTM | 78.02 | 72.85 | 82.47 | 74.16 | 0.5570 |
| pLSTM-CNN | 80.94 | 78.30 | 83.22 | 79.23 | 0.6164 |
| **SenSeqNet** | **83.55** | 79.53 | **87.03** | **81.92** | **0.6689** |

The CNN, known for its ability to capture spatial features, achieved an accuracy of 80.65% and an MCC of 0.6104. While effective in identifying local patterns within the protein sequences, the CNN struggled to capture temporal dependencies critical for detecting cellular senescence. To address this limitation, RNNs were introduced to model sequential data. The RNN showed an improvement, with an accuracy of 81.12% and an MCC of 0.6173, indicating its capability to better capture sequential patterns in the data. Next, more advanced temporal models, such as LSTMs, were explored. The LSTM, designed to handle long-term dependencies in sequential data, further improved accuracy to 82.09%, with an MCC of 0.6397. Building on this, the Bidirectional LSTM, which processes information in both forward and backward directions, enhanced the ability to capture complex dependencies, achieving an accuracy of 82.53% and an MCC of 0.6488. This model excelled at processing temporal patterns but lacked focus on spatial feature extraction.We also evaluated several hybrid architectures. Among these, the CNN-LSTM configuration demonstrated the lowest performance, with an accuracy of 78.02% and an MCC of 0.5570, suggesting that processing spatial features before sequential features may not fully leverage the strengths of both architectures. The pLSTM-CNN model performed better, achieving an accuracy of 80.94% and an MCC of 0.6164, though it had higher computational demands due to the simultaneous processing of spatial and sequential features.

The best-performing architecture was the SenSeqNet, which effectively combined the LSTM’s ability to capture sequence dependencies with the CNN’s proficiency in refining spatial features. SenSeqNet achieved the highest accuracy of 83.55%, an AUC of 0.92, and an MCC of 0.6689. These results demonstrate that the combination of LSTM and CNN architectures, when sequenced in this order, leads to a more comprehensive understanding of protein sequences, resulting in superior predictive accuracy. Overall, the results highlight that while single models such as CNNs and LSTMs are effective in their respective domains of spatial or temporal feature extraction, the hybrid approach of SenSeqNet provides the best of both worlds, outperforming both individual models and other hybrid configurations. The confusion matrix and ROC curve (Fig. 3E, F) further confirm the stability and generalizability of SenSeqNet in detecting cellular senescence.

### 3.4. Comparative analysis of SenSeqNet with machine learning models

To further evaluate the performance of SenSeqNet, we also compared its performance with various commonly used machine learning models including RF, XGBoost, SVM, and Bagging. The highest accuracy achieved among these machine models was 78.05% by RF, while XGBoost showed a notably high Sn of 87.67% but compromised Sp at 67.06%. Overall, none of the traditional models surpassed 80% in accuracy, with moderate MCC ranging between 0.5318 and 0.5583. The results are summaried in Table 4

**Table 4:** Comparative analysis of SenSeqNet with machine learning models on the independent test set.

| Model | Acc% | Sn% | Sp% | F1-Score% | MCC |
| --- | --- | --- | --- | --- | --- |
| RF | 78.05 | 77.11 | 79.37 | 80.39 | 0.5583 |
| XGBoost | 74.83 | **87.67** | 67.06 | 72.42 | 0.5318 |
| SVM | 77.02 | 79.01 | 74.79 | 78.38 | 0.5386 |
| Bagging | 77.98 | 77.51 | 78.60 | 80.14 | 0.5565 |
| **SenSeqNet** | **83.55** | 79.53 | **87.03** | **81.92** | **0.6689** |

In comparison, the SenSeqNet demonstrated superior performance, achieving accuracy of 83.55%, highlighting its effectiveness in extracting complex, non-linear features from large-scale protein sequence data. This disparity in performance underscores the capability of SenSeqNet to better handle the extensive and intricate relationships inherent in biological datasets, especially when dealing with large volumes of data, where traditional machine learning models may struggle to fully capture contextual dependencies.

### 3.5. Performance on Imbalanced Datasets

To assess the robustness of SenSeqNet in more realistic settings, we evaluated its performance on progressively imbalanced datasets with positive-to-negative ratios of 1:2, 1:3, 1:5, and 1:10, in addition to our original 1:1 dataset. For this updated analysis, we applied mechanisms specifically tailored to imbalanced learning, including focal loss^38^ and WeightedRandomSampler^39^, as well as threshold tuning^40^. These modifications are intended to bolster the detection of the minority class. Table 5 summarizes the results, reporting Accuracy (Acc), Sensitivity (Sn), Specificity (Sp), F1-score, and MCC under each ratio:

**Table 5:** Comparative performance of SenSeqNet on different imbalanced datasets.

| Ratios(P:N) | Acc% | Sn% | Sp% | F1-Score% | MCC |
| --- | --- | --- | --- | --- | --- |
| 1:1 | 83.55 | 79.53 | 87.03 | **81.92** | **0.6689** |
| 1:2 | 83.78 | **79.75** | 86.02 | 77.85 | 0.6511 |
| 1:3 | 85.60 | 75.90 | 88.98 | 73.12 | 0.6339 |
| 1:5 | 89.08 | 74.25 | 92.17 | 70.13 | 0.6362 |
| 1:10 | **93.25** | 72.87 | **95.29** | 66.25 | 0.6285 |

As the ratio of positives to negatives becomes increasingly skewed, we still observe a trade-off in classification metrics; however, the sensitivity (Sn) remains competitive. Notably, while overall accuracy escalates from 83.55% at 1:1 to 93.25% at 1:10, sensitivity only declines from 79.53% to 72.87%. Specificity concurrently rises from 87.03% to 95.29%, and we also observe modest gains in F1-score and MCC for several of the imbalanced ratios. These findings illustrate that applying class weighting, focal loss, and threshold tuning can substantially mitigate the risk of missing true positives in skewed datasets, allowing SenSeqNet to preserve more balanced performance across key metrics.

Nevertheless, the results show that as the class imbalance becomes more extreme, there remains a pronounced tendency to classify samples as “negative,” reflected in the continued (though improved) drop in sensitivity. In applications where missing a senescence-associated protein is especially costly, further refinements such as more extensive hyperparameter tuning or sophisticated oversampling/undersampling techniques may be warranted to further elevate Sn. By employing these tailored methods with minimal additional overhead, SenSeqNet demonstrates enhanced robustness in identifying senescence-associated proteins across various degrees of class imbalance.

### 3.6. External Validation for SenSeqNet

After incorporating specialized strategies (e.g., focal loss, WeightedRandomSampler, and threshold tuning) to address imbalanced data, we further validated SenSeqNet on an external dataset comprising 6,443 negatives and 2,750 positives (approximately a 1:2.5 ratio). These external samples were recently identified in 2024^41,42^ and were not included in any part of our model training or development. Despite this skew, the model continued to perform robustly, achieving a sensitivity of 78.18%, specificity of 80.82% and overall accuracy of 80.03%, with an F1 score of 70.08% and a Matthews correlation coefficient of 0.5601. The strong performance on these newly reported sequences confirms that SenSeqNet’s predictive ability is not merely an artifact of memorization but reflects underlying senescence-related features. This approach demonstrates how SenSeqNet can handle truly novel data—mirroring a real-world scenario in which newly discovered senescence-associated proteins must be classified without prior knowledge. Although the sensitivity is slightly lower than in our internal benchmarks, it remains sufficiently high for practical screening purposes, indicating the effectiveness of our imbalance-mitigation mechanisms in preserving the ability to detect true senescence-associated proteins without inflating false positives.

## 4. Conclusion

In this study, we have demonstrated the efficacy of a novel hybrid deep learning model SenSeqNet, which combining LSTM with CNN for the detection of cellular senescence from protein sequences. Our approach successfully leverages the sequential modeling capabilities of LSTMs and the spatial feature extraction strengths of CNNs to capture both temporal and spatial patterns within protein sequences that are indicative of cellular senescence. The SenSeqNet proved to be the optimal model, achieving an impressive accuracy of 83.55% and an AUC of 0.92 on the independent test set. The SenSeqNet outperformed both single models and other hybrid configurations, underscoring the importance of sequence processing order in deep learning architectures. To further evaluate SenSeqNet’s reliability, we performed 10-fold cross-validation, achieving a final cross-validated accuracy of 83.28% ± 0.58%, and validated the model on an external dataset, where it achieved an accuracy of 80.03%. These results confirm the robustness and generalizability of SenSeqNet.

Our findings highlight the potential of deep learning in advancing aging research by enabling accurate detection of aging at the cellular level. The robustness and scalability of the SenSeqNet model make it a valuable tool for large-scale screening in biological sequencing, providing rapid initial predictions of senescence-associated proteins. For instance, the model could be employed to identify aging-related proteins within the mitochondrial proteome, thereby offering valuable insights into mitochondrial dysfunction during aging. Furthermore, SenSeqNet could guide research into aging-associated diseases, such as Alzheimer’s disease and cardiovascular conditions, by pinpointing senescence-related proteins that may serve as potential biomarkers or therapeutic targets. Additionally, the model could contribute to drug discovery efforts by identifying key aging-related proteins for experimental validation and potential therapeutic intervention, particularly through the use of mouse models to test the efficacy of potential treatments. While it shows potential for identifying candidates for further study, its primary application lies in offering preliminary predictions and supporting subsequent validation efforts, rather than directly uncovering molecular mechanisms. This approach can significantly aid fundamental research by narrowing down targets for experimental investigation, ultimately contributing to a deeper understanding of cellular senescence.

However, a key limitation of this study is the model’s interpretability, particularly in identifying biologically significant motifs within protein sequences. Unlike extensively studied proteins, such as antibodies, where known motifs and experimental validation are available, proteins associated with cellular senescence lack such data. This gap limits our ability to confirm whether the model focuses on biologically meaningful patterns. Future research should address this challenge by exploring and validating potential motifs related to cellular senescence, enhancing both the interpretability and clinical applicability of the model’s predictions.

## Declarations

### Ethics approval and consent to participate

Not applicable.

### Consent for publication

Not applicable.

### Authors’ contributions

Hanli Jiang and Li Lin were responsible for the conceptualization and methodology development of the study, as well as drafting the original manuscript. Dongliang Deng and Siyi Liu contributed to data curation and investigation. Jianyu Ren and Xin Yang were involved in model development and visualization. Lubin Liu provided supervision, acquired funding, and led the review and editing of the manuscript. All authors have reviewed and approved the final version of the manuscript.

## Acknowledgements

We extend our sincere appreciation to the Program for Youth Innovation in Future Medicine at Chongqing Medical University for their valuable support.

## Availability of data and material

Publicly available datasets were analyzed in this study. The code utilized in this study is available at: https://github.com/HanliJiang13/SenSeqNet/.

